# Optogenetic control of Protein Kinase C-epsilon activity reveals its intrinsic signaling properties with spatiotemporal resolution

**DOI:** 10.1101/2025.01.06.631444

**Authors:** Qunxiang Ong, Crystal Jing Yi Lim, Yilie Liao, Justin Tze-Yang Ng, Ler Ting Rachel Lim, Shernys Xuan Yi Koh, Sher En Chan, Pheobe Lee Yu Ying, Huijun Lim, Chen Rui Ye, Loo Chien Wang, Siok Ghee Ler, Radoslaw M Sobota, Yaw Sing Tan, Gerald I. Shulman, Xiaoyong Yang, Weiping Han

## Abstract

The regulation of PKC epsilon (PKCε) and its downstream effects is still not fully understood, making it challenging to develop targeted therapies or interventions. A more precise tool that enables spatiotemporal control of PKCε activity is thus required. Here, we describe a photo-activatable optogenetic PKCε probe (Opto-PKCε) consisting of an engineered PKCε catalytic domain and a blue-light inducible dimerization domain. Molecular dynamics and AlphaFold simulations enable rationalization of the dark-light activity of the optogenetic probe. We first characterize the binding partners of Opto-PKCε, which are similar to those of PKCε. Subsequent validation of the Opto-PKCε tool is performed with phosphoproteome analysis, which reveals that only PKCε substrates are phosphorylated upon light activation. Opto-PKCε could be engineered for recruitment to specific subcellular locations. Activation of Opto-PKCε in isolated hepatocytes reveals its sustained activation at the plasma membrane is required for its phosphorylation of the insulin receptor at Thr1160. In addition, Opto-PKCε recruitment to the mitochondria results in its lowering of the spare respiratory capacity through phosphorylation of complex I NDUFS4. These results demonstrate that Opto-PKCε may have broad applications for the studies of PKCε signaling with high specificity and spatiotemporal resolution.

**Summary:** We have developed a photo-activatable optogenetic PKCε probe, which demonstrates differential activity in light *versus* dark. The tool is subsequently validated with protein association studies and phosphoproteome analysis. It enables dissection of signaling events arising from its activation at defined subcellular locations. For instance, sustained activation of PKCε at the plasma membrane is required for its phosphorylation of the insulin receptor at Thr1160.

## Introduction

PKCε is highly expressed in many cell types and involved in a diverse range of cellular processes, including cell proliferation, glucose metabolism, secretion, and memory formation. Consequently, dysfunctional PKCε activity is implicated in the pathogenesis of many diseases, including insulin resistance, type 2 diabetes and neurodegenerative diseases. PKCε is often linked to the formation of Aβ in Alzheimer’s Disease, increased cognitive deficits^1^ and reduced synaptogenesis^2^. The usage of PKCε activators such as DCP-LA and Bryotstatin-1 have been shown to reverse such effects^3,4^. PKCε has also been proposed as an oncogene and emerging tumor marker, which leads to the development and clinical trials for PKCε inhibitors^5^.

Much effort has been made to characterize the role of PKCε in cellular signaling. However, many challenges remain due to the highly complex nature of PKCε signaling. Firstly, PKCε is involved in compartmentalized cellular signaling that triggers site-specific behavioral phenotypes^6–8^. For instance, addition of phorbol ester 4β-phorbol-12-myristate-13-acetate (PMA) results in accumulation of PKCε at the plasma and nuclear membranes^9^. PKCε has also been reported to localize in Golgi apparatus^10^ and mitochondria^11^, modulating the functions of each organelle respectively. Secondly, PKCε is activated for varied periods of time in different circumstances^12,13^. The PKCs could undergo acute activation in the presence of GPCR or tyrosine kinase agonists, and a more sustained activation window is observed when agonists such as PMA are added^14^. Genetic overexpression and silencing experiments could also be affected by compensation effects. Most studies utilize chemicals, such as phorbol ester activators and inhibitors, for acute control of PKCε activity. However, these methods result in global changes in PKCɛ activity, and do not allow for compartmentalized dissection of PKCε signaling. In addition, off-target effects such as the simultaneous activation of other kinases and different PKC isoforms make it difficult to unravel PKCε signalling. Furthermore, since many of these inhibitors or activators cannot be metabolized, sustained modulation of PKCε occurs and results in poor temporal control^14^. Thus, obtaining a full grasp of PKCε signaling activities would require a more specific and precise method for dissection of specific pathways.

To overcome these hurdles, optogenetic systems are often used that allow for the precise spatiotemporal control of protein activation. The optogenetic systems have been used successfully to control the activity of kinases such as AKT1^15,16^, cRAF^17^, and TrkA^18^. In the present study, we have adopted optogenetics as a highly suitable approach for dissecting PKCε signaling cascades without the interference of other signaling nodes.

## Results

### Ideal Characteristics of Opto-PKCɛ

Engineering optical control of kinases requires the following considerations. Firstly, activation of opto-kinases should be controlled only by light without the need of exogenous factors. This will confer minimal dark background activation and full activation upon light illumination. Secondly, the downstream targets of the opto-kinase should resemble the kinase that it models. Thirdly, activation of the opto-kinase should be reversible and can be configured to activate at specific subcellular sites. In our study, we constructed an optically controllable PKCɛ that overcomes the limitations of current methods so as to precisely dissect PKCɛ signaling (**Figure 1A**).

**Figure 1:**
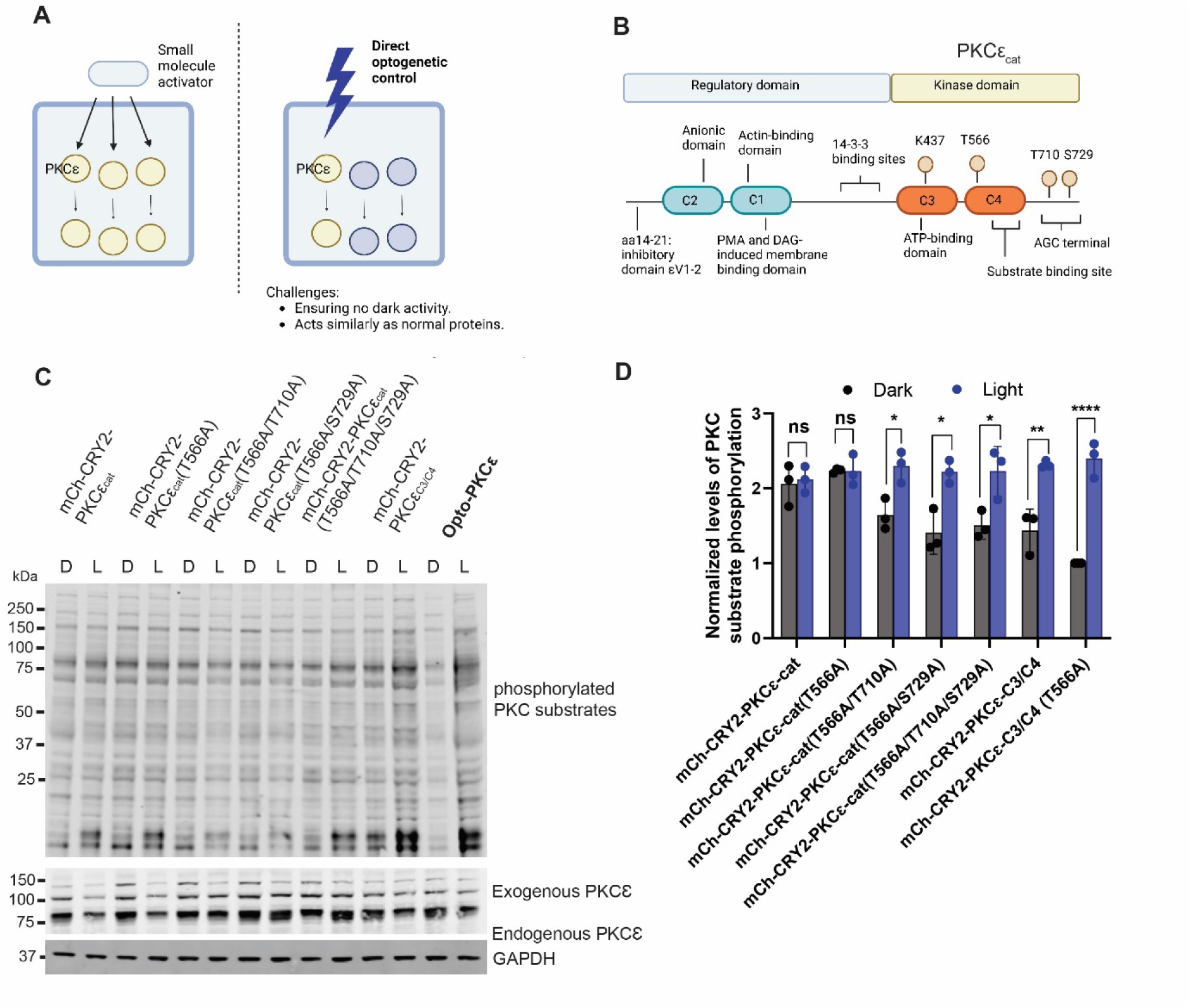
Design and development of Opto-PKCɛ. **(a)** Schematic of the ideal properties of an optically-controlled PKCɛ. Small molecule activators possess off-target effects, without spatiotemporal specificity. Direct optogenetic control of PKCɛ could allow dissection of the pathway if the challenges listed could be overcome. **(b)** Schematic of the regulatory and catalytic domains of PKCɛ. The regulatory domain of PKCɛ was truncated and replaced by the blue light sensitive cryptochrome domains tagged with mCherry fluorescent protein. **(c)** Representative Western blots of phosphorylation levels of PKCɛ substrates, PKCɛ and GAPDH for HEK293T cells transfected with constructs listed. These cells were either subjected to 5 minutes of dark condition or blue light. **(d)** Quantification (n=3) of normalized phosphorylation levels of PKCɛ substrates relative to GAPDH. The data is presented as mean +/− SD.

### Rational engineering of Opto-PKCɛ

PKCɛ activation involves the binding of membrane-resident diacylglycerol, which induces conformational changes and the release of the autoinhibitory pseudosubstrate site, resulting in the phosphorylation of three key signaling sites, Thr566, Thr710 and Ser729. (**Figure 1B**) Thr566 is phosphorylated by pyruvate dehydrogenase kinase 1 (PDK1) in a phosphoinositide-dependent manner, which leads to the autophosphorylation of two residues, Thr710 and Ser729 at the C-terminus^19^. Given that the first phosphorylation event initiated by PDK1 is canonically associated with the activation state of PKCɛ, it is critical to take into consideration the dependency of the phosphorylation sites for optimal optical control when designing the Opto-PKCɛ tool.

Considering that PKC stimulation results in self-association through intramolecular and intermolecular interactions between the regulatory and catalytic domains^20–22^, we tested if PKCɛ activation by PMA results in oligomerization of PKCɛ. We observed that mCherry-PKCɛ oligomerized after 5 minutes of PMA treatment (**Figure S1**). This result suggests light-induced oligomerization could be used to promote PKCɛ activation.

The endogenous PKCɛ consists of the regulatory domain (C2 and C1 domains) and the catalytic domain (C3, C4 domains and the AGC terminal). Referring to previous work on the phosphoinositide kinases^23^, Opto-Raf^24^ and Opto-AKT^25^, removal of the regulatory domain is typically needed to achieve full optical control of the optogenetic system. In place of the original regulatory domain, the mCherry-cryptochrome 2 (CRY2) construct was fused to the catalytic domain (**Figure 1B**). The CRY2 system has previously been known to promote oligomerization under blue light illumination.

Seven different constructs were tested in the optimization process and subjected to either dark or 15 minutes of blue light. Background PKCɛ activity was observed when the full catalytic domain was present while activity was reduced when Thr566, Thr710 and Ser729 were mutated to alanine. The difference in PKCɛ activity under dark and light conditions was further enhanced when the AGC terminal was removed. As such, we termed the fusion protein containing mCherry-CRY2 coupled with PKCɛ catalytic domain Ala566 and a truncated AGC terminal as Opto-PKCɛ (**Figure 1C**).

### Molecular dynamics simulations to understand the mechanisms of Opto-PKCɛ

To understand why the T566A mutation caused reduced kinase activity in the dark for Opto-PKCɛ, molecular dynamics (MD) simulations were performed to understand the conformational differences between T566-phosphorylated PKCε and the T566A mutant PKCε without the AGC terminal. As there were no available experimental structures of the catalytic domain of human PKCε, we used the AlphaFold model of human PKCε as the initial structure for the simulations^26^. This structure had the activation loop in an active conformation, where Thr566 was in close proximity with the positively charged residues Arg531 and Lys555. Conventional molecular dynamics (cMD) simulations were performed on the kinase domain of PKCε phosphorylated at Thr566 and the T566A mutant without the AGC terminal. Hydrogen bond occupancy analysis revealed that the negatively charged phosphate side chain of phosphorylated Thr566 (pThr566) forms intimate salt bridges with the positively charged side chains of Arg531 and Lys555, with average occupancies of more than 90% (97.9 ± 3.4 % and 90.1 ± 0.5 % respectively). These salt bridges stabilised the activation loop in the active conformation. When Thr566 was mutated to Ala, it did not interact with Arg531 and Lys555, which suggests that the active conformation of the activation loop could be unstable. The mutant PKCε showed slightly higher Cα root mean square fluctuations (RMSF) in a small region of the activation loop (residues 566–573) compared to phosphorylated PKCε (**Figure S2A**).

To determine if the activation loop underwent further conformational changes as a result of Thr566 phosphorylation and T566A mutation, we performed accelerated molecular dynamics (aMD) simulations to enhance conformational sampling. In these simulations, a boost potential was applied to lower the energy barriers for conformational transitions so that they could occur within a shorter timescale. The aMD simulations showed that the activation loop adopted significantly different conformations in the two systems (**Figure 2A, Figure S2B and C**). Compared to the cMD simulations, the dynamics of the phosphorylated PKCε was very similar, with the activation loop remaining in the active conformation and pThr566 in contact with Arg531 and Lys555, while the activation loop in mutant PKCε shifted towards the ATP binding site. In one aMD simulation run of mutant PKCε, this shifting of the activation loop was accompanied by an outward displacement of the αC helix (**Figure 2A**). This was likely to be an intermediate state between the active and inactive conformations of the activation loop. In the inactive conformation, the activation loop blocked the substrate from binding near the ATP binding site, thus preventing phosphorylation from occurring. These findings suggest that phosphorylation at Thr566 helps to stabilise the active conformation of the activation loop in PKCε and that the T566A mutant PKCε construct we used to form our Opto-PKCε tends towards the inactive conformation, thus accounting for the mutant construct’s reduced catalytic activity in the dark.

**Figure 2:**
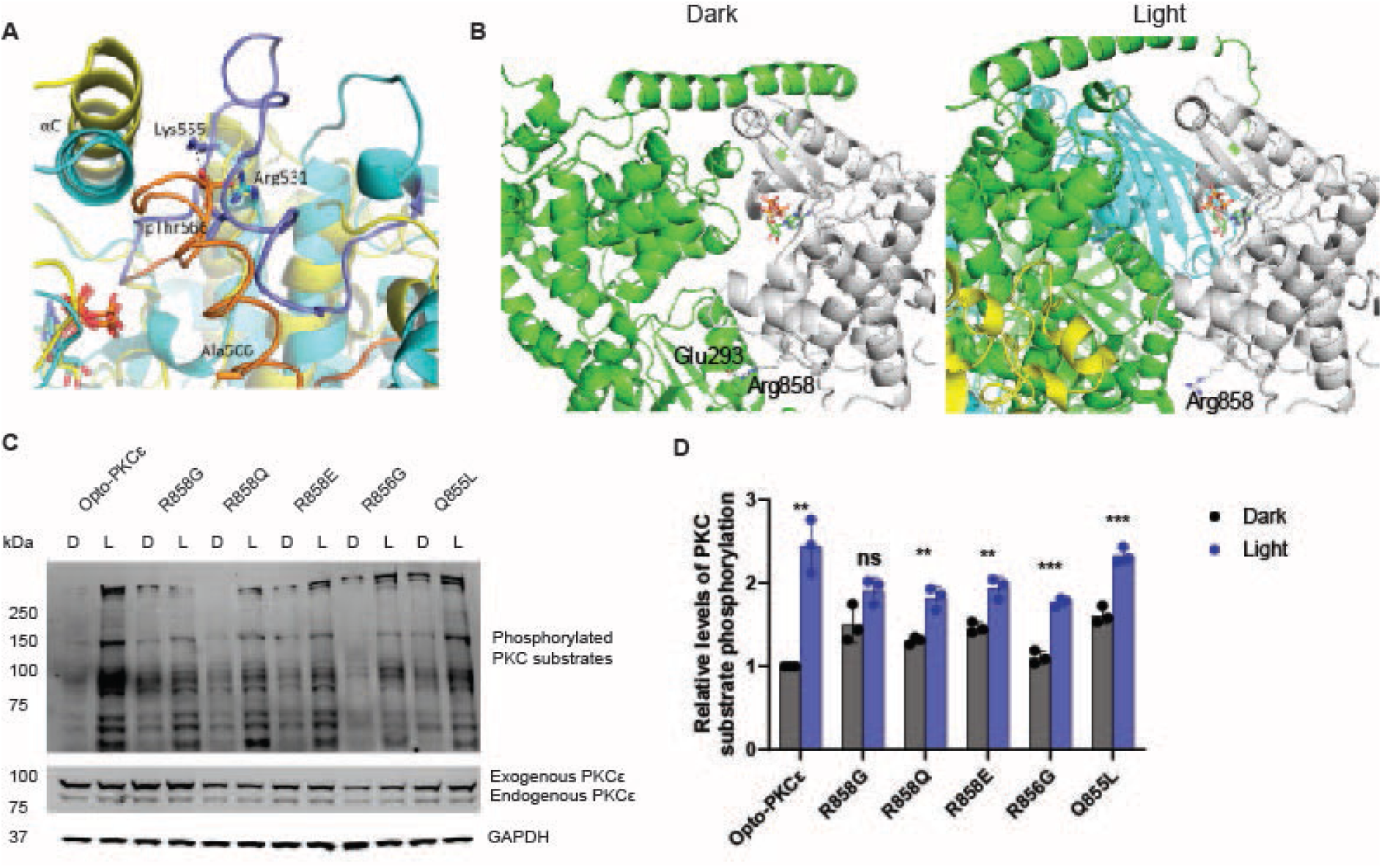
Computational simulations to aid in rationalization of Opto-PKCɛ dark-light activity. **(a)** Superposition of the final trajectory structures from representative aMD simulations of phosphorylated (cyan) and T566A mutant (yellow) PKCε. The activation loops of phosphorylated and T566A mutant PKCε were shown in purple and orange cartoon, respectively. **(b)** AlphaFold3 models of the monomeric (left) and tetrameric (right) Opto-PKCɛ reveal the Glu293-Arg858 interaction as a potential gatekeeper for dark-light activity. The PKCɛ domain is shown in white while the rest of the protein is shown in green. Other monomers in the tetramer are shown in yellow and cyan. Glu293, Arg858 and ATP are shown in sticks. **(c)** Representative Western blots of phosphorylation levels of PKCɛ substrates, PKCɛ and GAPDH for HEK293T cells transfected with constructs listed. These cells were either subjected to 5 minutes of dark condition or blue light. **(d)** Quantification (n=3) of normalized phosphorylation levels of PKCɛ substrates relative to GAPDH. The data is presented as mean +/− SD.

To determine the requirements for Opto-PKCɛ to be effectively light-gated, we examined the nature of the residues on the three key signaling sites. First, we compared Opto-PKCɛ with the construct containing phosphomimetic mutations (sites mutated to negatively-charged residues). We also compared constructs where Thr566 was mutated to glycine or valine, which are similar to alanine mutation. Opto-PKCɛ dead, which contains a K437R mutation that abrogates the ability to bind to ATP, was included as the negative control (**Figure S3A**). While the glycine and valine mutants showed similar light-dark properties as the Opto-PKCɛ, the triple negatively-charged mutant demonstrated higher levels of PKCɛ activity even in the dark state. This result implies that the charged activation at the signaling sites are crucial for mediating the catalytic domain’s activities, and the AGC terminal also affects PKCɛ activity control (**Figure S3B**).

Next, we examined how the positioning of the PKCɛ domain in the construct affects PKCɛ activation. We examined four constructs containing the optimized PKCɛ catalytic domain, namely mCherry-CRY2-PKCɛ (Opto-PKCɛ), CRY2-mCherry-PKCɛ, PKCɛ-CRY2-mCherry and PKCɛ-mCherry-CRY2. These constructs were subject to 15 minutes of darkness or blue light. Western blot analysis revealed that the activation of PKCɛ occurred only when the PKCɛ domain was fused at the C-terminus (**Figure S3C**).

### Structural modeling with AlphaFold3 to understand interactions gating dark-light activity of Opto-PKCɛ

We also seek to rationalize the gating of dark-light activity of Opto-PKCε. To this end, we used AlphaFold3 to predict structures of the CRY2 tetramer in light and its monomeric form in dark. The respective top-ranked models of the monomer and tetramer generated by AlphaFold3 suggest that tetramerization led to increased accessibility of the substrate binding site and ATP binding pocket, thus increasing kinase activity. In the monomer, the CRY2 and PKCε domains are in close contact, thus hindering substrate binding and restricting access to the ATP binding pocket. One of these interdomain interactions is the Glu293-Arg858 salt bridge, which is lost upon tetramerization. Nearby residues Arg856 and Gln855 may also be implicated. (**Figure 2B**) The loss of these interdomain interactions in the tetramer leads to the partial release of the PKCε domain from the CRY2 domain, thus increasing accessibility of the substrate and ATP binding sites, and consequently enzyme activity. To validate our hypothesis, we carried out mutations on Arg858, Arg856 or Gln855 and subjected these constructs to dark / light treatment. (**Figure 2C**) Mutation of Arg858 to glycine, glutamine (neutral) or glutamate (negative) consistently results in lower dynamic range of the dark-light activity, where PKC substrate phosphorylation level is higher in the dark and lower under blue light. (**Figure 2D**) This highlights the potential importance of residue Arg858 as a gatekeeper for the optical control of kinase activity. Mutation of Arg856 to glycine did not affect the PKC activity at dark, but results in lower PKC substrate phosphorylation level under blue light. All in all, the mutation analysis confirms the AlphaFold predictions where Arg858 and nearby residues play a key role in gating Opto-PKCε activity, where interactions such as Glu293-Arg858 may stabilize a conformation of Opto-PKCε that reduces accessibility of the ATP binding pocket to the substrate in the dark, and the absence of such interactions in light enables Opto-PKCε to phosphorylate substrates.

### Reversibility of Opto-PKCɛ

An advantage of intracellular optogenetic tools is the temporal control of the system. We subjected the cells transfected with Opto-PKCɛ to 15 minutes of blue light before 4 hours of darkness. The cells were then reactivated with 10 minutes of blue light. Western blot analysis revealed that the phosphorylation level of PKCɛ substrates remained high for the first 30 minutes before the substrates were dephosphorylated. Reactivation of the Opto-PKCɛ construct was achieved, thereby showcasing the reversibility of the system (**Figure S4**).

### Immuno-enrichment of Flag-Opto-PKCɛ and Flag-PKCɛ reveals similar substrate interactions

To confirm that Opto-PKCɛ functions similarly as endogenous PKCɛ, we next asked if the interactome of Opto-PKCɛ and PKCɛ are overlapping. We subjected HEK293T cells transfected with FLAG-tagged Opto-PKCɛ to five minutes of blue light and performed co-immunoprecipitation experiments to capture the proteins bound to the agarose FLAG beads (**Figure 3A**). Comparing the lists of proteins associated with FLAG-Opto-PKCɛ and FLAG-PKCɛ, we found that 238 out of 254 proteins are associated with both, while only 9 proteins each exclusively associate with each subset (**Figure 3B, Table S1**). The list of overlapping substrates includes PKCɛ, 14-3-3 proteins and MARCKS, which have been well validated in the past to be PKCɛ-associated proteins. These results reveal that the nature of the Opto-PKCɛ interactome to be similar to that of PKCɛ.

**Figure 3:**
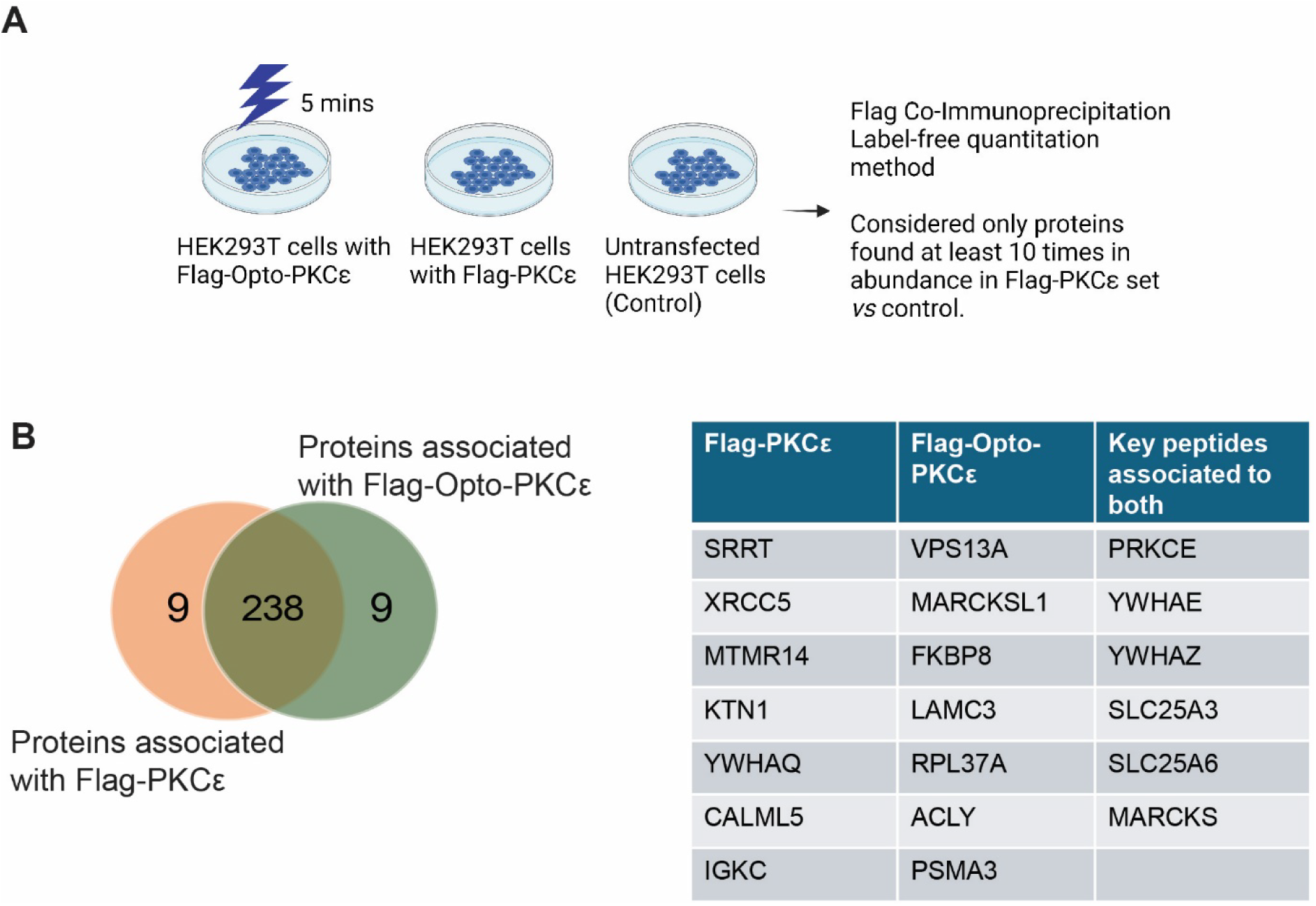
Opto-PKCɛ maintains largely similar interactome as that of PKCɛ. **(a)** Schematic of the workflow comparing interactome of Flag-Opto-PKCε and Flag-PKCε. Untransfected HEK293T cells was used as a negative control, with proteins found at least 10 times more in abundance considered for analysis. **(b)** Mass spectroscopy of proteins associated with Flag-Opto-PKCε or Flag-PKCε demonstrates that Flag-Opto-PKCε has a largely similar interactome as Flag-PKCε. Key proteins belonging to exclusively each subset or overlapping set is detailed.

### Comparison of signaling pathways activated by Opto-PKCɛ and PMA induction

Although previous studies have profiled cellular responses to PKCɛ activation by phorbol esters, the activators are also known to activate multiple pathways including PKA and other members of the PKC family. In contrast, Opto-PKCɛ allowed us to understand the direct downstream targets of PKCɛ exclusively (**Figure 4A**). To investigate the signaling pathways activated by Opto-PKCɛ and PMA, we subjected HEK293T cells to the following conditions. HEK293T cells expressing Opto-PKCɛ were stimulated with blue light for 15 minutes. In a separate set, HEK293T cells were exposed to PMA for 15 minutes.

**Figure 4:**
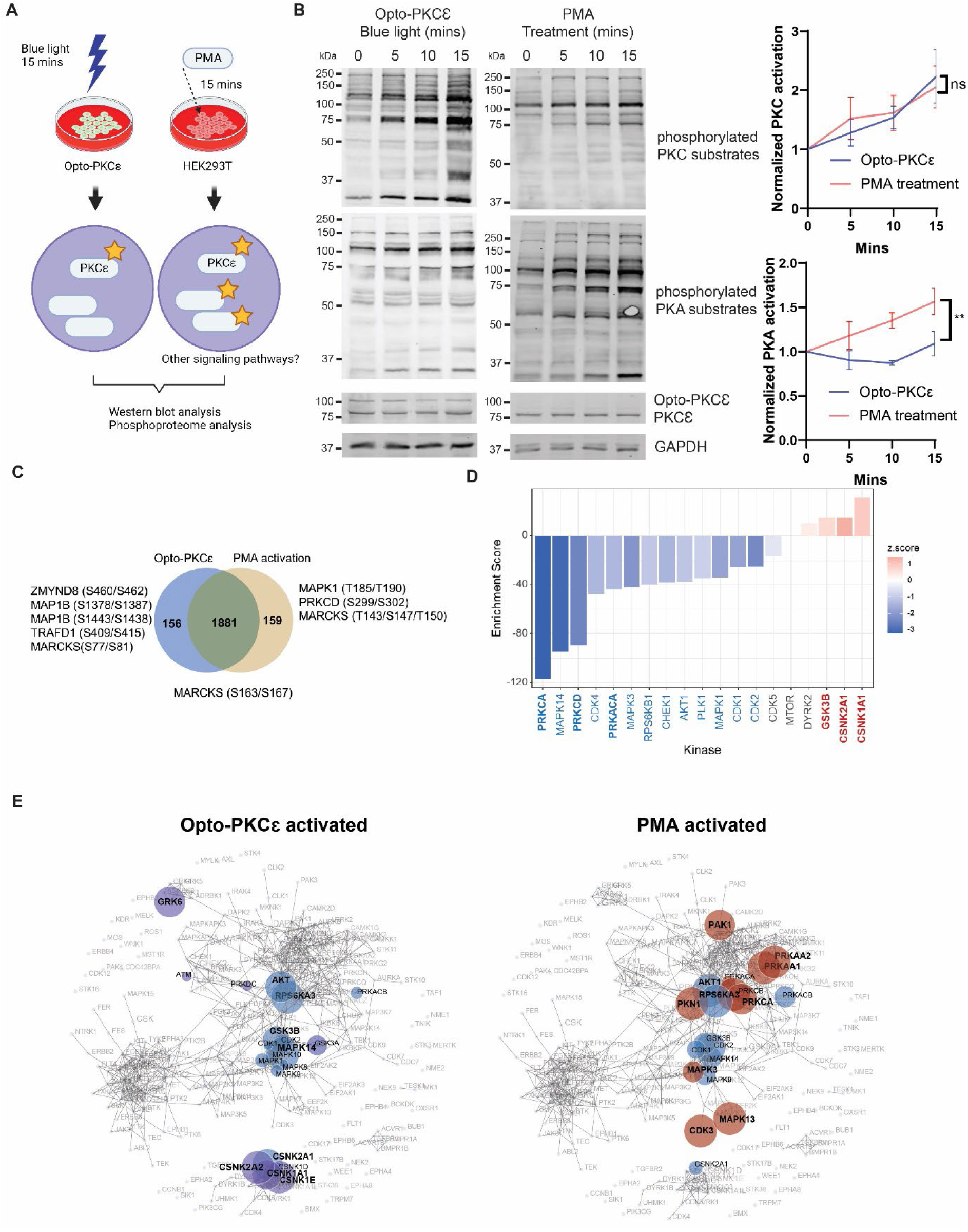
Opto-PKCɛ enables dissection of PKCɛ signaling. **(a)** Schematic of the experiments comparing signaling dynamics arising from 15 minutes of Opto-PKCɛ activation and PMA induction respectively. **(b)** Representative Western blots of phosphorylation levels of PKC and PKA substrates, PKCɛ and GAPDH for HEK293T cells subjected to both conditions. Quantification (n=3) of normalized phosphorylation levels of PKC and PKA substrates relative to GAPDH. **(c)** Venn Diagram of phosphopeptides quantifying the number of phosphopeptides appearing in both subsets or in either subset. Some of the main phosphopeptides known to be activated by PMA or classical PKCɛ phosphosites are highlighted. **(d)** KSEA revealed that CSNK and GSK3 substrates were enriched in opto-PKCε activation group while PKA pathways and other PKC pathways were enriched under PMA treatment. **(e)** KEA validated the increased phosphorylation of CSNK and GSK3 substrates under opto-PKCε activation (in purple) and those of PKA substrates (in red) under PMA treatment. Common substrates are highlighted in blue. The data in **(b)** are presented as mean +/− SD.

We first performed Western blot analysis on these cells to track their signaling patterns at 0, 5, 10 and 15 minutes of blue light exposure. As expected, both stimulation approaches resulted in an increase in phosphorylation levels of PKC substrates. However, only PMA caused PKA substrate phosphorylation (**Figure 4B**).

To broadly search for the downstream modules that could be activated by Opto-PKCɛ and PMA, we performed phosphoproteomics to have a comprehensive phosphorylation map. The lysates were collected and analyzed after stimulation. A comparison of the two conditions was detailed in **Table S2**. Out of 2196 phosphopeptides detected by phosphoproteomics, 159 phosphopeptides were found to be exclusively phosphorylated in the PMA treatment group, while 156 phosphopeptides were only observed in the Opto-PKCɛ activation group (**Figure 4C**). Ser163/Ser167 of MARCKS, well documented phosphorylation sites by PKCε, were indeed found to be phosphorylated under both conditions^27^. Notably, PKCɗ and MAPK1 were activated only by PMA and not Opto-PKCɛ, thereby suggesting that these substrates could be off-target effects attributed by PMA.

We then compared the peptides that had significant differences in phosphorylation levels in both conditions, as normalized to total protein levels (**Data S1**). Some of the more significant proteins were highlighted in the volcano plot in **Figure S5A**. 80 and 143 peptides were found to be more phosphorylated in Opto-PKCɛ and PMA conditions respectively for phosphopeptide quantification. We then conducted Kinase-Substrate Enrichment Analysis (KSEA) to estimate overall changes in a kinase’s activity based on the collective phosphorylation changes of its identified substrates^28^. We found that CSNK and GSK3 substrates were enriched in the Opto-PKCε activation group while PKA and other PKC pathways were enriched in the PMA treatment group (**Figure 4D**). We further validated this observation with Kinase Enrichment Assay (KEA), which allowed us to examine whether the gene sets from these two conditions were enriched with genes known to interact with kinases^29^. The results further confirmed the activation of pathways independent of PKCε-mediated pathways, such as PKA in the PMA-enriched subgroup (**Figure 4E**). Subsequently, we validated the increase in GSK3B and p38 phosphorylation in the Opto-PKCɛ subgroups (**Figure S5B and C**). On the other hand, the activation of MAPK1 was seen exclusively in the PMA subgroup (**Figure S5C**).

### Opto-PKCɛ can be customized to be activated at specific subcellular locations

While Opto-PKCɛ activation was seen in the entire cell, which was evident from the phosphorylation of proteins in all major subcellular compartments (**Table S3**), physiological activation of PKCɛ was usually site-specific. For instance, the translocation of PKCɛ to the plasma membrane has long been regarded as a readout for PKCɛ activation. In separate studies, PKCε has also been shown to be activated at the mitochondria, where PKCε phosphorylates the mitochondrial KATP channel in order to protect against ischemic insults^30,31^. To understand the activity of PKCɛ at specific subcellular locations, it is necessary to manipulate the selective recruitment of Opto-PKCɛ to the desired sites.

CRY2 forms heterodimers with crytochrome-interacting basic-helix-loop-helix 1 protein (CIB1) upon blue light stimulation. We made use of this feature to pair Opto-PKCɛ with fusion proteins of the N-terminus truncated version of CIB1 (aa 1-170), CIBN, tagged to cellular location markers^32^. Specifically, CIBN-GFP-miro and CIBN-GFP-CAAX were localized at the outer membrane of the mitochondria and plasma membrane respectively, allowing PKCɛ activation at these locations upon blue light recruitment (**Figure 5A**). Upon blue light stimulation, Opto-PKCɛ translocated to the plasma membrane in HEK293T cells co-transfected with Opto-PKCɛ and CIBN-GFP-CAAX (**Figure 5B**). Similarly, Opto-PKCɛ translocated to the mitochondria in HEK293T cells co-transfected with Opto-PKCɛ and CIBN-GFP-miro1 under blue light excitation (**Figure 5B**).

**Figure 5:**
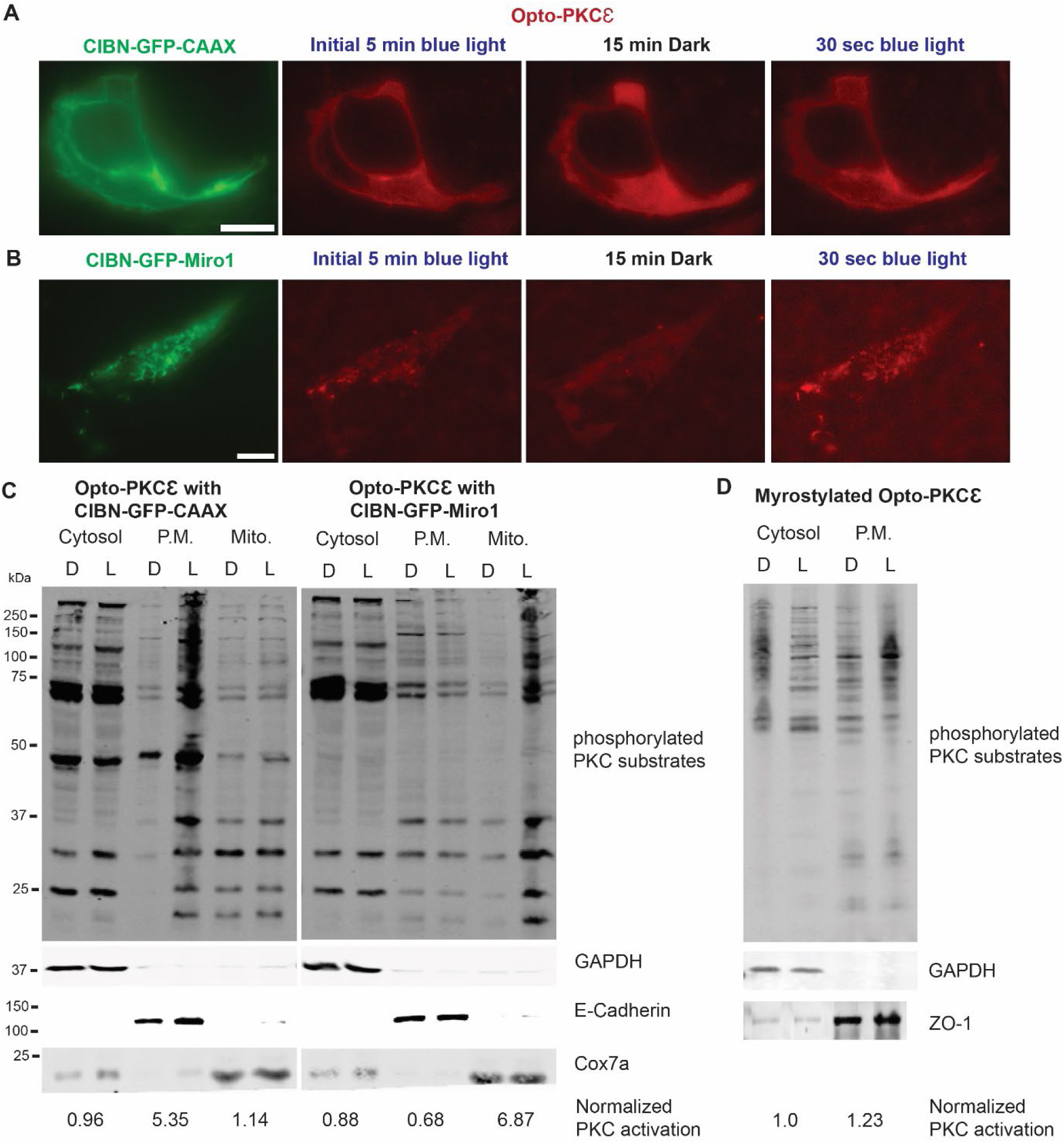
Opto-PKCɛ enables subcellular dissection of PKCɛ signaling. **(a)** Representative images of HEK293T cells transfected with Opto-PKCɛ and CIBN-GFP-CAAX from initial 5 min blue light to 15 minutes dark and then back to blue light exposure. CIBN-GFP-CAAX could be found to colocalize with Opto-PKCɛ after 30 seconds of blue light exposure. Scale bar = 10 µm **(b)** Representative images of HEK293T cells transfected with Opto-PKCɛ and CIBN-GFP-miro1 from initial 5 min blue light to 15 minutes dark and then back to blue light exposure. CIBN-GFP-miro1 could be found to colocalize with Opto-PKCɛ after 30 seconds of blue light exposure. Scale bar = 10 µm **(c)** Western blot analysis of subcellular fractionation experiment depicting phosphorylation of PKC substrates, E-cadherin, Cox7a and GAPDH of cells containing Opto-PKCɛ with CIBN-GFP-miro1 or CIBN-GFP-CAAX and subjected to 15 minutes of dark or blue light. **(d)** Western blot analysis of subcellular fractionation experiment depicting phosphorylation of PKC substrates, ZO-1 and GAPDH of cells containing myr-Opto-PKCɛ and subjected to 15 minutes of dark or blue light.

Using subcellular fractionation, we tested whether the recruitment of oligomerized Opto-PKCɛ to the different subcellular locations causes the local phosphorylation of PKC substrates. Indeed, when the HEK293T cells co-transfected with Opto-PKCɛ and CIBN-GFP-CAAX were exposed to blue light, phosphorylation of PKC substrates selectively increased only at the plasma membrane. When the localization tag was replaced with CIBN-GFP-miro, the increase in phosphorylation of PKC substrates occurred exclusively in the mitochondrial fraction (**Figure 5C**). Next, we tested whether the recruitment of Opto-PKCɛ from the cytosol is a necessary event, or if the same effect can be achieved if Opto-PKCɛ was already tagged to the desired localization. Interestingly, in the event where Opto-PKCɛ was constitutively localized at the plasma membrane through myristylation tag, the increase in phosphorylation of PKC substrates at the plasma membrane was less prominent (**Figure 5D**).

### Sustained PKCɛ activation at the plasma membrane of hepatocytes results in phosphorylation of the insulin receptor at Thr-1160

The role of hepatic PKCɛ in lipid-induced hepatic insulin resistance, particularly whether PKCɛ alone is responsible for phosphorylation of insulin receptor at Thr1160 in hepatocytes ^33,34,35^, has been under debate. Opto-PKCɛ tool is perfectly positioned to answer the question – (1) the use of the tool occurs in intact cells where only PKCɛ, and not other kinases, is selectively and precisely activated at the plasma membrane; (2) the duration of the activation can be tuned to investigate the nature of the interaction; (3) in contrast, drugs and genetic manipulation may cause compensatory effects, which may result in an indirect effect towards PKCɛ activation. We first checked the utility of the Opto-PKCɛ tool in isolated mouse hepatocytes and compared its activation in the cytosol versus the plasma membrane. Hepatocytes co-transfected with CIBN-GFP-CAAX and Opto-PKCɛ showed an increase in PKC substrate phosphorylation specifically at the plasma membrane (**Figure 6A**). Upon confirmation of the utility of Opto-PKCɛ tool in hepatocytes, we investigated whether PKCɛ activation alone at the plasma membrane would result in phosphorylation of insulin receptor at Thr-1160. Hepatocytes co-transfected with CIBN-GFP-CAAX and Opto-PKCɛ showed an increase in phosphorylation of insulin receptor at Thr1160 under blue light stimulation while this was not observed in hepatocytes transfected with only Opto-PKCɛ, highlighting the need for PKCɛ to be localized at the plasma membrane for the insulin receptor Thr1160 phosphorylation (**Figure 6B**). At the same time, this translated to a significant decrease in insulin-stimulated IRK Thr-1158/1162/1163 phosphorylation level for plasma membrane activation of PKCɛ. As a negative control, hepatocytes co-transfected with CIBN-GFP-CAAX and Opto-PKCɛ (dead) did not show any increase in phosphorylation of the insulin receptor at Thr-1160. In the absence of insulin, we noted an increase of the insulin receptor Thr1160 phosphorylation level as well (**Figure S6**). Lastly, we examined whether the Opto-PKCɛ needed to be constitutively localized at the plasma membrane for the insulin receptor Thr1160 phosphorylation. A transient bout of 15 minute Opto-PKCɛ activation at the plasma membrane was not sufficient to sustain the insulin receptor phosphorylation at Thr1160, indicating the temporal dependence of PKCɛ at the plasma membrane for the insulin receptor phosphorylation. (**Figure 6C**)

**Figure 6:**
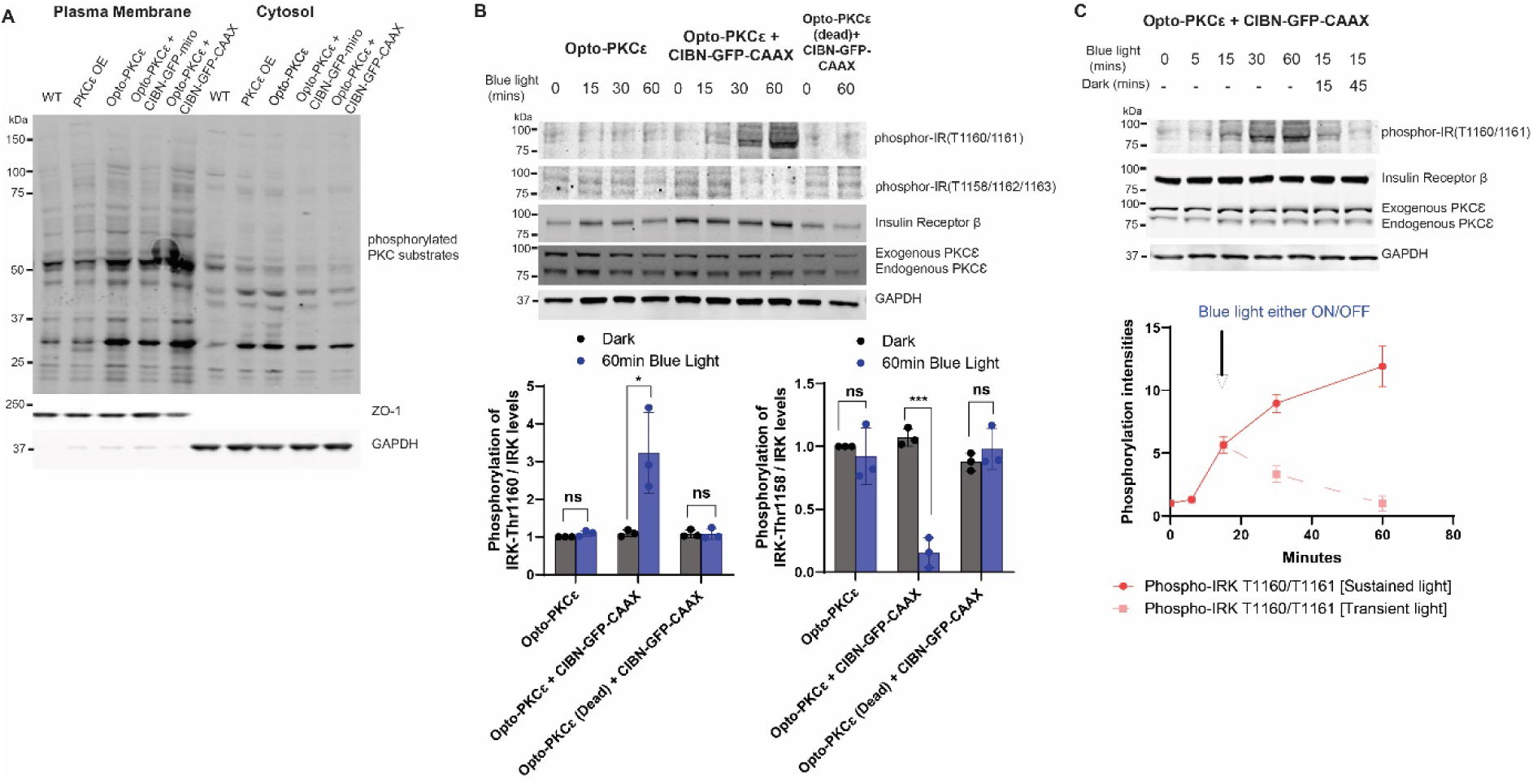
Sustained PKCɛ activation at the plasma membrane of hepatocytes results in phosphorylation of Insulin Receptor at Thr-1160. **(a)** Western blot analysis of subcellular fractionation experiment depicting phosphorylation of PKC substrates, ZO-1 and GAPDH of primary hepatocytes containing transfected plasmids as listed and subjected to 15 minutes of dark or blue light. **(b)** Representative Western blots of phosphorylation levels of phosphor-IRK (T1160/T1161), phosphor-IRK (T1158/T1162/T1163), Insulin Receptor, PKCɛ and GAPDH for primary hepatocytes subjected to stated conditions. Quantification (n=3) of normalized phosphorylation levels relative to insulin receptor levels. **(c)** Representative Western blots and phosphorylation levels of phosphor-IRK (T1160/T1161), Insulin Receptor, PKCɛ and GAPDH for primary hepatocytes subjected to stated conditions. Quantification (n=3) of phosphor-IRK (T1160/T1161) relative to Insulin Receptor levels. The data in **(b)** and **(c)** are presented as mean +/− SD.

### PKCɛ activation at the mitochondria results in reduced oxygen consumption rate

To investigate the role of PKCɛ activation at the mitochondria, we employed the Opto-PKCɛ tool coupled with CIBN-GFP-miro which was tagged at the outer mitochondrial membrane. We performed mitochondrial stress assays and measured the oxygen consumption rate (OCR) using different inhibitors to probe the bioenergetics parameters. **Figure 7A** showed the OCR of transfected HEK293T cells with or without blue light. Basal and ATP-dependent OCR readings indicate that PKCɛ activation at the mitochondria induced by light resulted in a lower OCR. We quantified the basal, ATP production, proton leak and spare capacities and found that light treatment significantly reduced these four parameters (**Figure 7B**). To ensure that this was not triggered by off-target effects due to blue light, we performed the mitochondrial stress assays on CIBN-GFP-miro-transfected cells with or without blue light, and did not observe any significant difference in the bioenergetics (**Figure 7C**).

**Figure 7:**
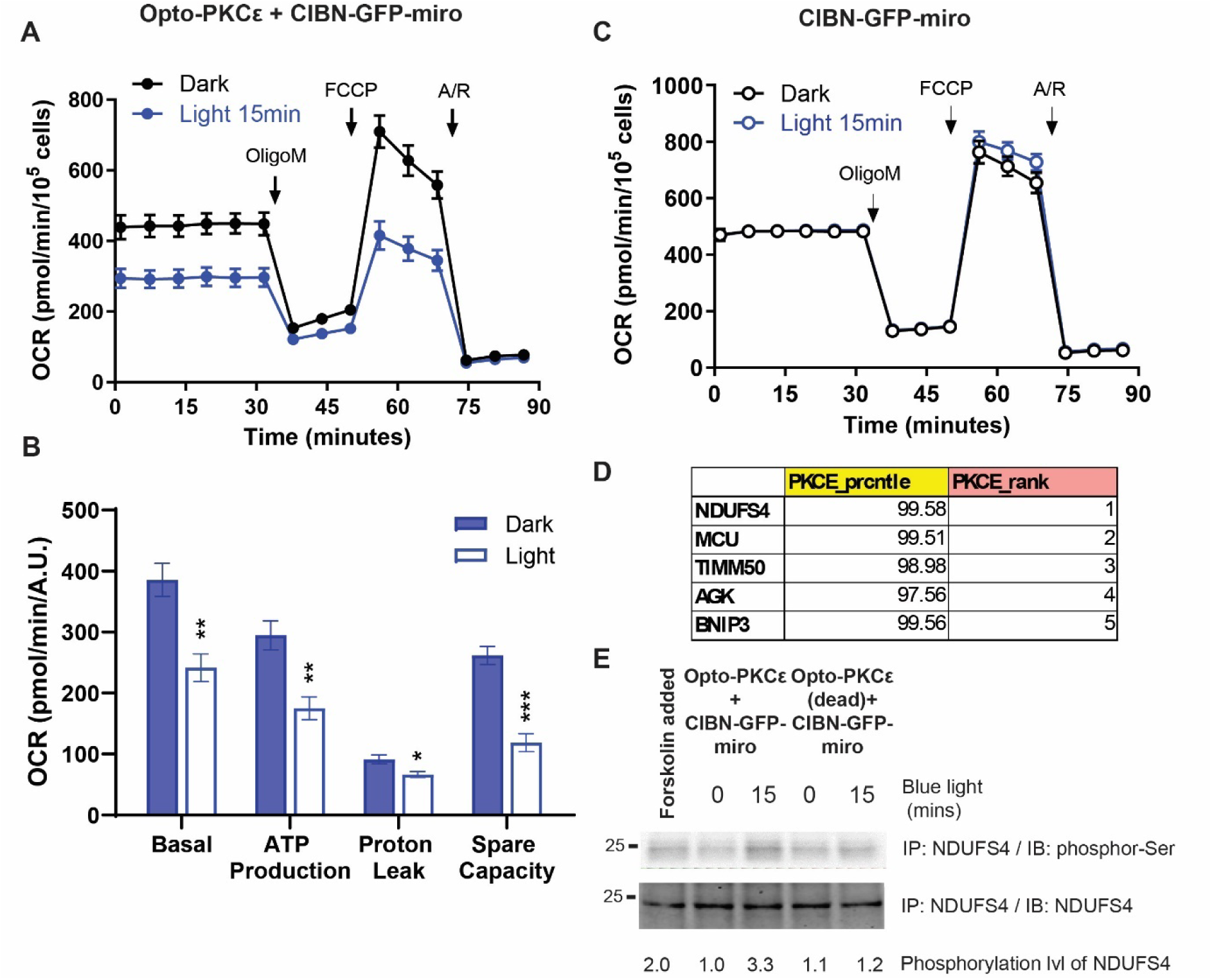
PKCɛ activation at the mitochondria results in reduced oxygen consumption rate. (a) HEK293T cells transfected with Opto-PKCɛ and CIBN-GFP-miro were subjected to 15 minutes of either dark or blue light. Oxygen Consumption Rate (OCR) measurements were then obtained with an extracellular flux analyser (Seahorse Bioscience). 4um of Oligomycin, 0.5um of FCCP, 1um of Antimycin A and Rotenone were injected into the cells at different time points preset on the machine and OCR response was recorded. 5 sets of experiments are represented. (b) Quantification (n=5) of the OCR of HEK293T cells transfected with Opto-PKCɛ and CIBN-GFP-miro under different conditions. (c) HEK293T cells transfected with CIBN-GFP-miro were subjected to either dark or 15 minutes of blue light. Oxygen Consumption Rate (OCR) measurements were then obtained with an extracellular flux analyser (Seahorse Bioscience). Similar inhibitor concentrations were used as in (a). (d) List of top phosphorylated substrates by PKCɛ that are known to be localized at mitochondria. (e) Western blots of co-immunoprecipitation experiments of NDUFS4 looking at phosphor-Ser and NDUFS4 levels subjected to stated conditions. Forskolin was added as a positive control. The data in **(a)** and **(b)** are presented as mean +/− SD.

To identify potential candidates associated with the bioenergetics phenotype, we listed the top phosphorylated mitochondrial substrates by PKCɛ as identified in a kinome-wide assay previously published (**Figure 7D**)^36^. NADH dehydrogenase [ubiquinone] iron-sulfur protein 4, mitochondrial (NDUFS4) was found to be the top candidate, which encodes a nuclear-encoded accessory subunit of the mitochondrial membrane respiratory chain NADH dehydrogenase (complex I). Indeed, phosphorylation of NDUFS4 was observed when PKCɛ activation at the mitochondria was induced by blue light (**Figure 7E**). This could explain the reduction in Complex I activity upon PKCɛ activation at the mitochondria^6^.

## Discussion

With more than 60,000 PubMed entries, PKC is one of the most well-studied kinases, and its many isozymes have been extensively characterized. However, it has been challenging to distinguish and disentangle the intricate signaling network of each PKC isozyme, and how the distinctive patterns caused by the activation of each PKC isozyme could lead to a vast range of cellular responses^37^. Due to the inherent limitations of traditional genetic tools and small molecule potentiators of PKC activity, the complexities of PKC activity and function have yet to be resolved.

Optogenetic control of individual PKC isozyme activity is an attractive strategy to address these fundamental questions. A typical approach in the field is to mimic the localization patterns of the intracellular proteins in its activation. This technique has been successfully utilized for the activation of PI3K^23^, Ras/Raf/ERK^24,38^, Bax^39^ and Rho-GTPases^40,41^. Indeed, a similar approach has been utilized by Logothetis and colleagues to recruit the catalytic domain of PKCɛ to the plasma membrane^42^; the activation of GIRK1/4 was the readout for PKCɛ activation. However, it is important to note that PKCɛ has been shown to exert signaling effects on multiple subcellular locations, such as Golgi^10^, mitochondria^30,31^ and nucleus^43^. As such, it is important to ensure the background activity of opto-PKCɛ is effectively eliminated, and that the presence of blue light would be the only trigger required to activate the ideal opto-PKCɛ construct. This is the first report of an PKCɛ optogenetic tool with reduced background activation that may potentially affect the cellular signaling dynamics.

To achieve minimal background activation of the optogenetic construct, we took reference from previous work on the activation of optoPKA.^44^ In that study, mutation of residues at the activation loop allowed the construct to have dark-silent and light-active properties. Hence, we developed an optogenetic strategy that allows us to achieve minimal activity in the dark state. To do so, we inactivated the phosphorylation site Thr566 at the activation loop by mutating the site to an alanine. This is a key requirement to realize the robust light-dark differentiation of Opto-PKCɛ. We then demonstrated that the removal of the AGC terminal further differentiated the activity of Opto-PKCɛ under blue light. The AGC terminal is a key regulatory element of protein kinases A, G and C, which help to structure the catalytic core and promote the stability of PKCɛ through phosphorylation of the turn motif and hydrophobic motif^45^. In our study, we found that the removal of the AGC terminal did not affect the stability of Opto-PKCɛ but might lessen restriction of the catalytic core, thus allowing activation upon light-induced oligomerization.

We then tested whether Opto-PKCɛ behaves similarly to endogenous PKCɛ. Our phospho-proteomics analysis showed that the substrates phosphorylated by Opto-PKCɛ overlaps greatly with those phosphorylated by endogenous PKC, which affirms that the Opto-PKCɛ tool does mimic the activity of endogenous PKC. Our phosphoproteomics analysis has also provided further insights into the substrates of PKCɛ. Notably, MAPK1 was found to be activated only by PMA and not Opto-PKCɛ. Classical overexpression studies have suggested that Raf, an upstream kinase of MAPK1, and PKCɛ activate each other^46^. However, a study has demonstrated that PKCɛ does not activate Raf-1 and could be indirectly stimulating Raf1 by inducing the production of autocrine growth factors^47^. Next, we found that the activation of Opto-PKCɛ resulted in the activation of p38 pathway, a phenomenon not being observed through PMA induction. Furthermore, previous work has highlighted that the PKCɛ-p38 axis^48^, and its activation could have been potentially masked by other pathways, such as PKA and other PKC isoforms, that are also activated by PMA. Opto-PKCɛ activation is specific and avoids activating kinase-mediated pathways that can mask signaling pathways regulated by PKCɛ.

Opto-PKCɛ also potentially paves the way in understanding phosphosites arising exclusively from PKCɛ activation, which could be masked by the activation of other signaling pathways. For instance, while MARCKS was activated on the known canonical site (S163/167) by both PMA and Opto-PKCɛ, S77/S81 phosphorylation was only observed in the Opto-PKCɛ set while T143/S147/T150 phosphorylation was only observed in the PMA subset. In addition, MAP1B, also known to be phosphorylated by PKCɛ^49^, has additional sites that were phosphorylated by the Opto-PKCɛ not found in the PMA treatment group. As the functions of the S77/S81 and T143/S147/T150 phosphosites on MARCKS and the phospho-sites on MAP1B are virtually unknown, Opto-PKCɛ can be a useful tool to unravel the role of these phospho-sites.

In this paper, we have attempted to understand the potential mechanisms driving the dark-light activity of Opto-PKCɛ. While the oligomerization activity of PKCɛ upon PMA induction has not been reported before, there have been reports on PKCɑ oligomerization^20–22^. As such it is not surprising to observe the PKCɛ oligomerization events due to the higher concentration of PKCɛ on lipid rafts and potential transphosphorylation activities which would require close proximity^50^. Interestingly, light-induced oligomerization of Raf1 has also shown to induce its activation^17^. Nonetheless, the forced oligomerization of PKCɛ without the regulatory domain may induce conformational changes in the accessibility of the ATP binding site, which could be a future area of investigation for the design of optogenetic kinases. Alphafold simulations suggest that Glu293-Arg858 intramolecular interactions might play a role in reducing the accessibility of the kinase catalytic domain in the dark state, and the absence of such interaction in the oligomerized light state enables its activity to be reconstituted. The role of Glu293 (Glu42 of cryptochrome) was not included in this study as we hypothesized that the crptochrome oligomerization activity may be affected if mutations were to be performed here. As such, the mutation studies are focused on the residues on PKCɛ (R502, R500, Q499 corresponding to R858, R856 and Q499 on Opto-PKCɛ), which are not located immediately at the ATP binding site. The phosphorylation levels at both dark and light levels were within the original dynamic range of Opto-PKCɛ activity, suggesting that mutations on these residues did not affect the intrinsic kinase activity.

Unlike previous studies where kinase domains are recruited to plasma membrane to elicit their activity^25,42^, the Opto-PKCɛ tool is not reliant on translocation activity and overcomes the limitations to probe the role of PKCɛ only on the plasma membrane. Achieving subcellular targeting and probing its downstream signaling are now possible in other subcellular compartments. We also attempted to create a myrostylated form of Opto-PKCɛ where changes in PKC substrate phosphorylation were not as dramatic when compared to recruitment of Opto-PKCɛ to the plasma membrane via cotransfection of CIBN-GFP-CAAX and Opto-PKCɛ. This could be due to the localized high concentration of the myrostylated Opto-PKCɛ, which may contribute to higher background activity. Further work should focus on fine-tuning the expression levels in attempt to reduce the background activity. In addition, Opto-PKCɛ sets the stage to study specific phosphorylation events on a PKCɛ substrate, and such concepts can be extended to other PKC isozymes to understand the signaling dynamics by each individual isozyme.

Previously, PKCɛ has been shown to be activated by increased plasma membrane associated *sn*-1,2 diacylglycerol levels in hepatic insulin resistance^51^. While we did not observe the landmark event of IRK Thr1160 phosphorylation by both Opto-PKCɛ and PMA^33,34^ in HEK293T cells, we carried out the activation of Opto-PKCɛ in mouse hepatocytes where we subjected the transfected cells with different spatiotemporal windows of activation. Indeed, the sustained localization of activated Opto-PKCɛ is required for the IRK Thr1160 phosphorylation, while insulin-stimulated IRK Thr1158/1162/1163 phosphorylation is downregulated. While the Thr1160 phosphorylation event has been convincingly demonstrated in the past in hepatocytes overexpressed with constitutively membrane-localized PKCɛ and also *in vitro* settings, the experiments performed with the optical PKCɛ present direct evidence for Thr1160 phosphorylation of IRK by PKCɛ only if PKCɛ is localized at the plasma membrane, and if its activation is sustained.

The advent of the tool also allowed us to probe the role of PKCɛ at other organelles. Traditionally, it has been assumed that PKCɛ activation is noted by its translocation to the plasma membrane, but some reports have noticed the translocation of PKCɛ to other locations^6^. For instance, oxidant-induced injury causes PKCɛ to activate and translocate to the mitochondria in kidney cells and that this activation mediates decreases in mitochondrial function, specifically in complex I-mediated respiration, active Na+ transport, and Na+-K+-ATPase activity^6,52^. Our results support these findings and suggest a potential role of NDUFS4 phosphorylartion by PKCε in mediating the observed phenomena. More studies specifically at finding the site of phosphorylation on NDUFS4 could be carried out for further exploration.

The Opto-PKCɛ tool could easily be applied to other prevailing questions in biology. For instance, the role of PKCɛ can be dissected in cancers where different PKC isozymes can have similar, and sometime opposing roles in cell proliferation and other cellular functions^53^. Moreover, the role of PKCɛ activation in memory formation has yet been elucidated. This could be addressed with the application of Opto-PKCɛ in rodent brains^54^.

### Limitations of the study

The Opto-PKCɛ tool had not been studied for the number of reversible cycles and longer durations to be utilized. More studies could be performed to ascertain the underlying reasons that govern the dark-light activity of Opto-PKCɛ, for example cryo-EM crystallography could enable the structural features to be studied in detail. Additionally, our optical system requires the use of overexpression systems, so the system may not be easily applicable to cell lines where transfection may be difficult. It is also possible that in our FLAG pulldown experiments that some nonspecific proteins may still be identified despite having negative controls. Lastly, we have not yet applied the tool to tissues in an animal model; however, we foresee that the deployment of the components using AAVs could ensure targeting to particular tissues without need for additional modifications.

## Data availability statement

The data are available from the corresponding author upon reasonable request

## Acknowledgements

This work was supported by the A*STAR Biomedical Research Council core fund, A*STAR Strategic Research Program (the Brain-Body Initiative, iGrants call ID #21718), A*STAR Use-Inspired Basic Research Award to W.H. Q.O. gratefully acknowledges support from the NMRC (National Medical Research Council) YIRG grant (OFYIRG19nov-0045). X.Y. was supported by grants from National Institutes of Health (R01DK089098, R01DK102648, R01DK137467). G.I.S. was supported by grants from the US Public Health Services NIH/NIDDK (R01 DK133143 and P30 DK045735). J.T.N. and Y.S.T. are supported by the Bioinformatics Institute, A*STAR and A*STAR’s BMRC Central Research Fund award. We would like to thank Dr. Xiao-Bing Gao and Prof Bianxiao Cui for their feedback and comments.

## Author contributions

Conceptualization - Q.O.

Methodology - Q.O., R.M.S., Y.S.T., G.I.S. and W.H.

Data curation and investigation - Q.O., C.J.Y.L., L.T.R.L., S.K., S.E.C., P.L.Y.Y., H.L., J.T.N., L.C.W. and S.G.L.

Formal analysis - Q.O., C.J.Y.L., Y.L., J.T.N. and Y.S.T. analyzed the data.

Writing original draft – Q.O. and C.L.

Writing: review and editing - Q.O., Y.S.T, X.Y. and W.H.

## Supplemental Information

Document S1. Figures S1-S6 and Table S4.

Table S1-S3. Excel file containing data too large to fit into a PDF.

Data S1. Excel files and graph analysis files containing data too large to fit into a PDF.

## Key resources table

| REAGENT or RESOURCE | SOURCE | IDENTIFIER |
| --- | --- | --- |
| **Antibodies** |  |  |
| PKC epsilon monoclonal anti-Rabbit | Cell Signalling Technology | #2683 |
| Phospho-(Ser) PKC substrate polyclonal anti-Rabbit | Cell Signalling Technology | #2261 |
| Insulin Rb antibody (C-19) polyclonal anti-rabbit | Santa Cruz Biotechnology | sc-57342 |
| Phospho-IRK (T1150/T1151) monoclonal anti-mouse | Sigma Aldrich | 04-299 |
| Phospho-IRK (T1163/1162/1158) polyclonal anti-rabbit | Sigma Aldrich | 07-841 |
| Phospho-IRK (T1160) monoclonal anti-rabbit | Novo Nordisk |  |
| P44/42 MAPK polyclonal anti-rabbit | Cell Signalling Technology | #9102 |
| Phospho-p44/42 MAPK (Thr202/Tyr204) polyclonal anti-rabbit | Cell Signalling Technology | #9101 |
| P38 MAPK (D13E1) XPI monoclonal anti-rabbit | Cell Signalling Technology | #8690 |
| Phospho-p38 MAPK (Thr180/Tyr182) polyclonal anti rabbit | Cell Signalling Technology | #9211 |
| GSK3b | Cell Signalling Technology | #9315 |
| Phospho-GSK3b (Ser9) | Cell Signalling Technology | #5558S |
| GAPDH monoclonal anti-mouse | Santa Cruz Biotechnology | sc-32233 |
| NDUFS4 monoclonal anti-mouse | ThermoFisher | MA5-19432 |
| Phospho-Serine | Abcam | Ab9332 |
| E-Cadherin (HECD-1) | ThermoFisher | #13-1700 |
| VDAC | Abcam | ab34726 |
| IRDye® 800CW Goat anti-Rabbit IgG (H + L) | LICOR | 926-32211 |
| IRDye® 680RD Donkey anti-Mouse IgG (H + L) | LICOR | 926-68070 |
| Alexa Fluor 488 Goat anti-rabbit | ThermoFisher | A11008 |
| 4',6-Diamidino-2-Phenylindole, Dihydrochloride (DAPI) | ThermoFisher | D1306 |
| **Bacterial and virus strains** |  |  |
| One Shot Stbl3 Comp E Coli | Invitrogen | C737303 |
| **Biological samples** |  |  |
| Mouse: C57BL/6 | InVivos | N/A |
| **Chemicals, peptides, and recombinant proteins** |  |  |
| Phorbol 12-myristate 13-acetate (PMA) | MedChemExpress | HY-18739 |
| Forskolin PKA Activator | Sigma-Aldrich | F3917 |
| PKCε Inhibitor Epsilon-V1-2 (ε-V1-2) | MedChemExpress | HY-P0154 |
| Fetal bovine serum (FBS) | Gibco | 10500064 |
| Horse Serum | Thermofisher Scientific | 16050122 |
| Penicillin-Streptomycin (P/S) | Gibco | 15140122 |
| Dulbecco's Modified Eagle Medium High Glucose | Cytiva | SH30243.FS |
| M199 media | Gibco | 11150067 |
| Seahorse XF DMEM Media | Agilent | 103680-100 |
| Trypsin | Gibco | 12605036 |
| Poly-D-Lysine | Gibco | A3890401 |
| ExpiFectamine™ 293 Transfection Kit | Gibco | A14525 |
| Opti-MEM™ I Reduced Serum Medium | Gibco | 51985091 |
| PBS, pH 7.4 | Gibco | 10010023 |
| Collagenase Type IV | Sigma Aldrich | C5138 |
| EGTA | EMD Millipore Corp. | 324626 |
| c0mplete, EDTA-free EASYpack | Roche | 4693159001 |
| PhosSTOP™ EASYpack | Roche | 4906845001 |
| Triton-X100 | Sigma Aldrich | 9036-19-5 |
| Tween-20 | Promega | H5152 |
| 4X LDS | Invitrogen | NP0008 |
| Bovine Serum Albumin (BSA) | Capricorn scientific | BSA-1S |
| 10X Tris-Glycine-SDS buffer | BioRad | 1610772 |
| 10X Tris-Glycine buffer | BioRad | 1610771 |
| 20X MOPS SDS Running buffer | Invitrogen | NP0001 |
| 20X Tris Buffered Saline Buffer | Base Asia | BUF-3030-20X4L |
| *Dimethyl sulfoxide* (DMSO) | Sigma Aldrich | D8418 |
| In-Fusion® Snap Assembly Master Mix | Takara USA Bio | 638944 |
| FavorPrep Plasmid Extraction Kit | Favorgen Biotech | FAPDE300 |
| PureLink™ HiPure Plasmid Filter Maxiprep Kit | Invitrogen | K210016 |
| Phusion Flash High Fidelity PCR MasterMix | ThermoFisher | F548L |
| Critical commercial assays |  |  |
| Minute™ Plasma Membrane Protein Isolation and Cell Fractionation Kit | Invent Biotechnologies | SM-005 |
| Agilent Seahorse XF Cell Mito Stress Test Assay | Agilent | 103015-100 |
| Pierce™ BCA Assay | ThermoFisher | 23250 |
| Pierce™ Protein A/G Magnetic Agarose Beads | ThermoFisher | 78610 |
| Experimental models: Cell lines |  |  |
| Human: Human Embryonic Kidney cells HEK293T (female) | ATCC | CRL-3216 |
| Human: HepG2 (Male) | ATCC | HB-8065 |
| Animal: African green monkey kidney fibroblast-like cell-line COS7 | ATCC | CRL-1651 |
| Oligonucleotides |  |  |
| Primers for Opto-PKC tool, see **Table S4** | Present study | N/A |
| Software and algorithms |  |  |
| ImageJ | National Institute of Health | N/A |
| GraphPad Prism | GraphPad Software Inc., La Jolla, CA, USA | N/A |
| Kinase-Substrate Enrichment App (KSEAapp) | R Studio | N/A |
| Recombinant DNA |  |  |
| CIBN-GFP-CAAX | Addgene | #26867 |
| CRY2high-mCherry | Addgene | #104063 |
| Flag-PKCε | Addgene | #10795 |
| Other |  |  |
| LED light array | ThorLabs | LED465E |
| XF24 Extracellular Flux Analyzer | Seahorse Biosciences | 102238-100 |

## METHODS AND MATERIALS

### Cell culture

Human embryonic kidney 293T (HEK293T) and Human hepatoblastoma HepG2 cells were cultured in Dulbecco’s Modified Eagle Medium High Glucose (DMEM HG) supplemented with 10% Fetal Bovine Serum (FBS) and 1% Penicillin-streptomycin (P/S).

PC12 cells were cultured in F12-K media supplemented with 15% horse serum and 2.5%FBS. All cell plates were maintained in a standard incubator at 37°C in an atmosphere of 5% CO_2_.

### Primary mouse hepatocytes

Hepatocytes were isolated from C57BL/6 female mice between 8-12 weeks old. Perfusion buffer containing 1mm EGTA and 5.5mm glucose in HBSS was infused into the liver via the portal vein for 10 mins using a 25G butterfly needle at a speed of 3ml/min. Digestion buffer containing 0.3mg/ml of Collagenase Type TV, 5mm CaCl2 and 5.5mm glucose was thereafter introduced into the liver for 10 mins at a speed of 3ml/min. The liver was subsequently harvested and dissociated with attachment media containing 0.2% Bovine Serum Albumin, 2% Fetal Bovine Serum and 1% Penicillin streptomycin in media M199 media using forceps. Cell suspensions were passed through two 70 μm strainers per liver and underwent centrifugation twice at 50g for 2 mins. The cells were re-suspended and washed in attachment media after each centrifugation. Cell count and viability were performed using trypan blue and an automated cell counter. Tissue culture dishes were pre-coated with 0.1% fish gelatin for 1h and washed twice with PBS (Gibco). Live cells were plated at a density of 10 million per 10cm. Cells were subsequently incubated in a 37°C incubator containing 5% CO2 for 4 hours before attachment was aspirated and replaced with maintenance media containing only M199. Subsequent experiments performed on the isolated hepatocytes were done within 48 hours of isolation.

### Proteomics sample preparation (on beads digestion-LC-MS/MS)

Samples were reconstituted with 50 μL of 100mM Triethylammonium bicarbonate (TEAB), followed by adding 50 μL of 2,2,2-Trifluoroethanol (TFE). Samples were then reduced by adding to the final concentration of 10 mM of Tris(2-carboxyethyl) phosphine (TCEP) and incubated at 55°C for 20 minutes shaking at 900 rpm on a thermomixer. The samples were cool down to room temperature before speedvac to remove 2,2,2-Trifluoroethanol (TFE). Around 20 μL of the samples were left and alkylated to final 55mM of 2-Chloroacetamide (CAA) in dark for 30 minutes. All samples were top up to 100 μL with 100mM TEAB and digested with 4 μg of LysC for 4 hours, followed by 4 μg of trypsin overnight at 37°C with thermomixer at 900 rpm.

Desalting was carried out using HLB C18 reversed-phase cartridges. Desalted peptides were reconstituted with 10 μL of 0.1% (v/v) formic acid in water and 2 μL was injected for mass spectrometry analysis. Chromatography separation was performed using Easy-nLC 1200 (Thermo Fisher Scientific) coupled to an Orbitrap Fusion Lumos mass spectrometer (Thermo Fisher Scientific) using data-dependent mode. Peptides were separated using a 2-30% (v/v) acetonitrile gradient over 80 min, increasing to 45% over the next 5 min, and finally to 95% over 5 min, running at a constant flow rate of 300 nl/min.

The following parameters were set for MS data acquisition: automatic gain control (AGC) customed at normalized target of 100% with orbitrap resolution at 60,000 ranging from m/z 350 to 1550. Dynamic exclusion for precursors selected for fragmentation was set to 90 s. For fragmentation, MS2 isolation window was set to 1.2 m/z of the selected precursor masses and collision-induced dissociation (CID) was activated at 35% normalized collision energy. Fragment signals (MS2) were analysed by an ion trap analyzer set to normal mode, AGC target customed at 150% and maximum injection time (IT) at dynamic mode.

Raw mass spectra were searched using Sequest available in Proteome Discoverer™ (v3.1, Thermo Fisher Scientific) against human primary protein sequences retrieved from uniprot (19 July 2022) with 42289 sequences and 24370425 residues. The mass tolerance for precursors and fragments were set to 20 ppm and 0.8 Da, respectively; and relative quantitation was based on label-free quantitation (LFQ) method.

### Phosphoproteomics

Cell pellets were washed with phosphate-buffered saline (PBS) twice and supernatant was removed completely. Samples were then resuspended in 8 M urea in 50 mM 4-(2-hydroxyethyl)-1-piperazineethanesulfonic acid (HEPES), pH 8.0 and sonicated on ice using a probe sonicator at 30-s pulse, 30-s pause for 90 s. Total protein amount was estimated using bicinchoninic acid (BCA) assay and 3 mg proteins from each sample were aliquoted into new tubes. Aliquoted proteins were reduced 10 mM final concentration of tris(2-carboxyethyl)phosphine (TCEP) and incubated for 20 min at 55 °C for disulfide bridge reduction followed by alkylation with 55 mM 2-chloroacetamide (CAA) in the dark for 30 min. The concentration of urea in each sample was reduced to 1 M using 100 mM triethylammonium bicarbonate (TEAB), pH 8.5, then digested with endoproteinase LysC (1:40 enzyme:protein ratio) for 3 h and subsequently by trypsin (1:25 enzyme:protein ratio) at 37 °C overnight. Digestion was terminated by adding 1% (v/v) final concentration of trifluoroacetic acid (TFA) to the samples, followed by desalting using Sep-Pak C18 reversed-phase cartridges. Desalted peptides were separated, where 100 µg of peptides from each sample was taken for whole proteome analysis while the remaining was lyophilized. Lyophilized peptides were resuspended in 6 ml solvent mixture containing 40% (v/v) acetonitrile (ACN) and 6% trifluoroacetic acid (TFA). Phosphopeptide enrichment was carried out by loading the resuspended peptides to 10 mg of titanium dioxide (TiO_2_) beads pre-activated using dihydrobenzoic acid. Incubation was performed on a rotator at 25 °C for 10 min. The samples were centrifuged using 1000 × g for 1 min and the supernatant was transferred to a new tube containing 10 mg of TiO_2_ beads. Following incubation and centrifugation as described above, the beads were combined in new tubes and washed sequentially with the following buffers: 6% TFA in 10% ACN once; 6% TFA in 40% ACN twice; and finally 6% TFA in 60% ACN twice. Samples were dried by vacuum centrifugation to remove all residual volumes of ACN and TFA.

Phosphopeptides were eluted from the beads by sequential addition and collection of the following buffers: 5% ammonium hydroxide (after 15 min of incubation on a shaker at 25 °C) and 25% ACN in 10% ammonium hydroxide. For each buffer, centrifugation at 14,000 × g was carried out to pellet down the beads and the supernatant was transferred to new tube. Both eluates were combined as a single sample and filtered through a C8 membrane to remove any beads. The membrane was washed with 100% ACN to elute any bound peptides, combined with the initial eluate, and dried by vacuum centrifugation. Dried peptides were washed once using 60% CAN containing 0.1% (v/v) formic acid and re-dried in preparation for mass spectrometry analysis.

Dried fractions were resuspended in 10 µl of 2% (v/v) ACN containing 0.06% (v/v) TFA and 0.5% (v/v) acetic acid and transferred to an autosampler plate. Online chromatography was performed in a Vanquish Neo (Thermo Fisher Scientific) liquid chromatography system using a single-column setup and 0.1% formic acid in water and 0.1% formic acid in 80% acetonitrile asmobile phases. Fractions were injected and separated on a reversed-phase C18 analytical column (Easy-Spray, 75 µm inner diameter × 50 cm length, 2 µm particle size, Thermo Fisher Scientific) maintained at 50 °C and using a 0-35% (v/v) acetonitrile gradient over 60 min, followed by an increase to 50% over the next 8 min, and to 100% over 2 min. The final mixture was maintained on the column for 5 min to elute all remaining peptides. Total run duration for each sample was 75 min at a constant flow rate of 300 nl/min except for the isocratic hold at the final 5 min at 400 nl/min.

Data were acquired using an Orbitrap Eclipse mass spectrometer (Thermo Fisher Scientific) using data-dependent mode. Samples were ionized using 2.1 kV and 300 °C at the nanospray source. Positively-charged precursor signals (MS1) were detected using an Orbitrap analyzer set to 60,000 resolution, automatic gain control (AGC) target of 400,000 ions, and maximum injection time (IT) of 50 ms. Precursors with charges 2-7 within an isolation window of 1.2 m/z of the selected precursor masses and having the highest ion counts in each MS1 scan were further fragmented using collision-induced dissociation (CID) at 35% normalized collision energy. Fragment signals (MS2) were analysed by an ion trap analyzer set to Rapid mode, AGC target of 10,000 and maximum IT of 35 ms. Precursors used for MS2 scans were excluded for 60 s to avoid re-sampling of high abundance peptides. The MS1–MS2 cycles were repeated every 3 s until completion of the run. Proteins were identified using Proteome Discoverer™ (v3.0, Thermo Fisher Scientific). Raw mass spectra were searched using Sequest against human primary protein sequences retrieved from Swiss-Prot (11 June 2019). Carbamidomethylation on Cys were set as fixed modification; deamidation of asparagine and glutamine, acetylation on protein N-termini, phosphorylation of Ser, Thr, and Tyr residues, and methionine oxidation were set as dynamic modifications for the search. Trypsin/P was set as the digestion enzyme and was allowed up to two missed cleavage sites. Precursors and fragments were accepted if they had a mass error within 20 ppm and 0.8 Da, respectively. Peptides were matched to spectra at a false discovery rate (FDR) of 1% (strict) and5% (relaxed) against the decoy database and quantitated using label-free quantitation (LFQ) method. Search result was exported and further analyzed as described below.

### Phosphoproteomics Analysis

Phosphopeptides with at least 2 detectable values out of 3 replicates in at least one group were used for subsequent normalization. Peptides containing valid observations in fewer than 2 replicates were considered as undetected in all 3 samples within this group. The intensity of each phosphopeptide was normalized to the abundance of the corresponding protein. To identify differentially regulated phosphopeptides, missing values were replaced with half of the minimum peptide abundance within each samples, then log 2 (fold changes) and p-values were generated by comparing Opto-PKCɛ activated group with PMA treated cells. Phosphopeptides with |log 2 (fold changes)| ≥ 1 and p-value > 0.05, or those only detected in 1 group, were considered as specific phosphorylation events in either Opto-PKCɛ activation or PMA treatment group. Kinase-Substrate Enrichment Analysis (KSEA) was performed by the R package KSEAapp using default cutoff settings and the annotations from both PhosphoSitePlus and NetworKIN. Kinase Enrichment Analysis (KEA) was conducted by KEA2 webtool.

### Molecular Modelling

#### System setup

The model of human PKCɛ was obtained from the AlphaFold database^26^. The initial structure for the simulations comprised residues 408–737, which corresponds to the protein kinase domain (residues 408–668) and the C-terminal tail (residues 669-737). A protein BLAST search was conducted on Protein Data Bank (PDB) structures using the FASTA sequence of the region included in our model to identify similar PDB structures containing the ATP ligand and its two associated Mg^2+^ ions bound at the ATP binding site, since they were absent in the AlphaFold model. The highest ranked structure in terms of E-value was the crystal structure of mouse PKA catalytic domain in complex with phospholamban (PDB ID 7E0Z^55^). This structure was aligned to the AlphaFold model of human PKCɛ before transferring over the ANP and Mg^2+^ to the latter. To convert the ANP to ATP, the nitrogen atom between the beta and gamma phosphate was converted to an oxygen atom. Two systems were set up for simulation, one being T566A mutant PKCɛ with no C-terminal tail (residues 408–668) and the other being PKCɛ phosphorylated at Thr566 (residues 408–737). For the system with phosphorylated threonine, the coordinates of phosphorylated threonine were modelled using PyMOL^56^.

For each system, the N-terminus of the protein chain was capped with an acetyl group. PDB2PQR^57^ was used to determine residue protonation states and add hydrogen atoms. Using the LEaP module of AMBER 22, each system was solvated with TIP3P^58^ water molecules in a periodic truncated octahedron box such that the distance between the protein and box edge was at least 10 Å. Sodium counterions were then added to neutralize the systems.

#### Molecular dynamics (MD) simulations

Energy minimization and MD simulations were performed by the PMEMD module of AMBER22^59^ using the ff14SB^60^ field for the protein and the generalized AMBER force field (GAFF)^61^ for the ATP. Parameters for phosphorylated threonine and ATP were used as described by previous works^62,63^. Bonds involving hydrogen atoms were constrained using the SHAKE^64^ algorithm to enable a time step of 2 fs. A nonbonded interaction cutoff distance of 9 Å was used. Long-range electrostatic interactions were treated using the particle mesh Ewald^65^ method under periodic boundary conditions. Energy minimization was performed for 500 steps with the steepest descent algorithm, followed by another 500 steps with the conjugate gradient algorithm. The system was then heated gradually to 300 K over 50 ps in the NVT ensemble before equilibration in the NPT ensemble at a constant pressure of 1 atm for another 50 ps. Weak harmonic positional restraints with a force constant of 2.0 kcal mol^−1^ Å^−2^ were imposed on heavy atoms during energy minimization and equilibration steps. Further equilibration without positional restraints was performed for 2 ns, followed by the production run of 500 ns at 300 K and 1 atm. For each system, three simulation runs of conventional molecular dynamics (cMD) simulations with differing initial velocities were performed. The temperature of the system was maintained using a Langevin thermostat^66^ with a collision frequency of 2 ps^−1^, while the pressure was maintained using the Berendsen barostat^67^ with a pressure relaxation time of 2 ps.

#### Accelerated molecular dynamics (aMD) simulations

In aMD, a boost potential ΔV(r) is added to a system’s potential energy when the system’s potential energy is lower than the threshold energy value E.

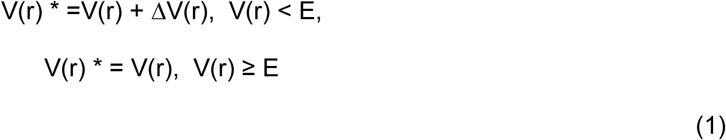

In this equation above, V(r)* is the modified potential energy, V(r) is the unmodified original potential energy, and ΔV(r) is the boost potential. In the dual-boost implementation of aMD, a boost potential is applied to the system’s potential energy and dihedral energy as follows:

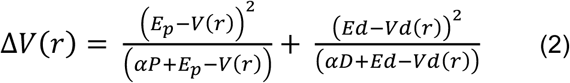

In the equation above, E_p_ is the threshold potential energy, E_d_ is the threshold dihedral energy, V(r) is the system’s potential energy, and Vd(r) is the system’s dihedral energy. αP is the acceleration factor for the system’s potential energy and αD is the acceleration factor for the system’s dihedral energy. The boost parameters were _calculated_ as follow:

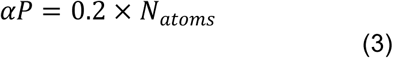

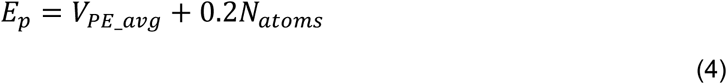

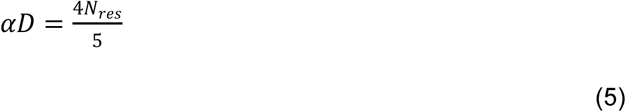

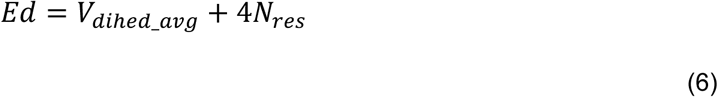

In the above equations, N_atoms_ is the total number of atoms, N_res_ is the number of protein residues, V_PE_avg_ is the average potential energy and V_dihed_avg_ is the average dihedral energy. V_PE_avg_ and V_dihed_avg_ for aMD were calculated for each system as the average value obtained from the corresponding 500-ns cMD simulation runs. Dual-boost aMD simulations, in which both dihedral energy and total potential energy are boosted, were initiated from the final structures of each corresponding cMD simulation runs for each system. All aMD simulations were performed at 300 K and 1 atm. Three runs of aMD simulations were performed on each system for 500 ns.

#### Trajectory analysis

Analysis was performed using the CPPTRAJ^68^ module in AMBER22. Hydrogen bond occupancy was calculated using the hbond command. The criteria for hydrogen bond formation are: an acceptor-heavy atom–donor-heavy-atom distance of less than 3.5 Å, and an acceptor heavy atom–donor hydrogen atom–donor heavy atom angle of more than 135⁰. Hydrogen bond occupancy analysis was performed on the last 400 ns of each simulation run to account for initial equilibration.

### AlphaFold3 modeling

Models of monomeric and tetrameric Opto-PKCɛ were generated using the AlphaFold Server (http://alphafoldserver.com), which is based on the AlphaFold3 model^69^. To generate models of the monomer, an amino acid sequence corresponding to Opto-PKCɛ, 2 ATPs and 1 FAD were used as input whereas for the tetramer, copies of Opto-PKCɛ, 8 ATPs and 4 FADs were used as input..

### Plasmid cloning

CIBN-GFP-CAAX; CRY2high-mCherry (Plasmid #104063) and Flag-PKCɛ plasmids were purchased from Addgene. For all other plasmids mentioned in this paper, the list of primers is recorded in **Table S3**. All PCR reactions were done using PhusionFlash High Fidelity Polymerase, followed by Gibson Assembly or InFusion. All plasmids were grown in Stbl3 bacteria and extracted using FavorPrep Plasmid Extraction Kit. Sequences were confirmed using BioBasic Asia Pacific Sequencing Services. Subsequently, amplification and purification were done using Invitrogen™ PureLink™ HiPure Plasmid Filter Maxiprep Kit. Gel-electrophoresis was done to ensure that all synthesized plasmids were clean and of the correct size. Amplified plasmids were stored at −20°C freezer and used for transfection.

### Plasmid transient transfection

The day before transfection, HEK293T and HepG2 were split to a density of 2.5*10^6^ cells in 10cm culture dishes and grown in DMEM HG + 10% FBS without P/S media. Cells were transfected using the Expifectamine 293 transfection kit and protocol. For each plate, 20ug of plasmid, 54ul of Expifectamine reagent and 2ml of Opti-MEM were used. No enhancer reagents were added the following day and no media change was carried out unless high cell death was observed. Other than using m199 media instead of DMEM HG, transfection for primary mouse hepatocytes was done in the same way.

### Light-delivery system

For light illumination experiments, a 3-by-2 blue LED array was constructed as described in adapted from an earlier report^24^. The array comprises of 24 blue LEDs (LED450L, Thorlabs). The light intensity at the height (12 cm) where the cell culture plate is placed was measured by a power meter (Newark, 1931-C) to be at 200 µW/cm2 with an absolute measurement uncertainty of less than 5%. The working distance of 12-cm is to ensure uniform blue light intensity across the surface of the cell culture dishes.

### Optogenetic experiments

Transfected cells were exposed to blue illumination (200 µW/cm^2^) using a homemade LED light array while dark plates were kept in the incubator wrapped in aluminium foil. Cell plates exposed to blue light are lysed under ambient light while dark plates are lysed with the aid of red light. For dark recovery, the cells were exposed to blue light before being wrapped in aluminium foil and placed back in the 37°C incubator.

Cells were washed twice with PBS before lysing with 400ul of NP40 lysis buffer supplemented with cOmplete protease inhibitor and PhosSTOP as per manufacturer’s instructions.

### Plasma membrane and Mitochondria Isolation

Subcellular fractionation of transfected HEK293T and hepatocytes was performed using the Minute™ Plasma Membrane Protein Isolation and Cell Fractionation Kit (Invent Biotechnologies). Steps were followed according to the manual provided.

### Co-immunoprecipitation Assay

After cell lysis and protein BCA Assay quantification, samples were diluted to a concentration of 500ug/ml in 500ul using NP40 lysis buffer. Subsequently, 5ul of respective primary antibody was added into each tube and rotated at 4°C overnight before co-immunoprecipitation was carried out as written in the manufacturer’s protocol handbook.

### Western blot

After lysed samples were agitated at 4°C for 1 hour and centrifuged at 16000g for 20 minutes, supernatant was obtained and quantified using Protein BCA assay on Tecan plate reader. 4X LDS with β-mercaptoethanol buffer was then added to samples and heated at 95°C for 5 minutes before subjecting to electrophoresis using Bio-Rad’s MiniPROTEAN system. After separation, protein lanes were transferred onto a nitrocellulose membrane using wet transfer at a constant 300mA for 90 minutes. Membrane was subsequently washed with 0.1% TBS-Triton (TBST) before standard blocking procedure. Primary antibodies used were listed in **Key Resource Table.**

### Fluorescence Microscopy – Live Cell Imaging

35mm glass bottom dishes were coated with 250ul of poly-D-Lysine for 1 hour before rinsing thrice with autoclaved water. COS7 cells were then plated at a density of 0.8*10^5^ cells for each dish in DMEM HG media + 10% FBS the day before transfection. 2.5ug of plasmid was transfected using Expifectamine reagent and Opti-MEM as mentioned above and imaging was carried out 24 hours after transfection. To observe the reversibility of Opto-PKCɛ, images were taken every minute under TRITC filter (557nm, 200ms pulse duration). On the other hand, images were taken every 5 seconds while rotating between FITC (∼495nm, 50ms pulse duration) and TRITC (557nm, 200ms pulse duration) filter to observe Opto-PKCɛ activation. Cells were not continuously shone under the FITC filter to prevent photobleaching. Images were obtained with Nikon Eclipse Ti-E inverted microscope (Nikon Imaging Centre, Singapore), with a CoolSNAP HQ2 CCD Camera, under 100x oil-immersion objective lens (N.A. 1.49).

### Immunostaining

COS7 cells were plated at a density of 0.8*10^5^ on a 35mm poly-D-lysine coated glass bottom dish and transfected with Opto-PKCɛ using Expifectamine transfection protocol as mentioned above. Opto-PKCɛ plate was either treated with continuous blue light for 15 minutes or wrapped with aluminium foil before fixation process. Cells were washed thrice with PBS followed by fixing with 4% PFA for 10 minutes at room temperature. Following that, cells were washed thrice with PBS and incubated in permeabilization solution for 5 minutes. This is followed by blocking using 10% FBS in PBS for 1.5 hours in a 37°C incubator. Cells were then stained with primary antibodies for 1 hour. After washing thrice with PBS, cells were then incubated with secondary antibody (Alexa Fluor 488, 1:300) and wrapped in aluminium foil for 1 hour. Detailed information regarding 1^st^ and 2^nd^ antibodies was provided in the **Key Resources Table**. Images were taken using a conventional fluorescence microscope with an oil-immersion 100x objective lens.

### Seahorse XF Cell Mito Stress Assay

Reagents and apparatus used to perform Seahorse XF Cell Mito Stress Test Kit were all purchased from Agilent. 24 well seahorse culture plates were first coated with poly-D-lysine and incubated in 37°C incubator containing 5% CO2 for 1 hour before washing twice with PBS. Rows C and D were wrapped with aluminium foil and secured with aluminium tape. Plate covers were also covered in aluminium tape to prevent exposure of blue light to the cells. Plates and covers were treated with UV for 30 mins in the culture hood before transfected HEK293T cells were seeded at 0.4*10^5^ cells per well in DMEM HG + 10%FBS + 1% P/S. 18 hours after seeding, cells were incubated in the Seahorse XF DMEM Medium supplemented with 10Mm glucose and 2mm sodium pyruvate for 45mins in a 37°C incubator with no CO2. Media change was performed in the dark. Cells were subsequently exposed to 15mins of blue light and loaded into the seahorse machine probe. Probes were incubated in calibrant overnight before the experiment. 4um of Oligomycin, 0.5um of FCCP, 1um of Antimycin A and Rotenone were injected into the cells at different time points preset on the machine and OCR response was recorded.

### Quantification

All quantifications were done using Image J, Graph analysis was done using GraphPad Prism Software 8.4.3.

